# An artificial intelligence-based first-line defence against COVID-19: digitally screening citizens for risks via a chatbot

**DOI:** 10.1101/2020.03.25.008805

**Authors:** Alistair Martin, Jama Nateqi, Stefanie Gruarin, Nicolas Munsch, Isselmou Abdarahmane, Bernhard Knapp

**Author notes:** these authors contributed equally to this work (Alistair Martin, Jama Nateqi).

## Abstract

To combat the pandemic of the coronavirus disease (COVID-19), numerous governments have established phone hotlines to prescreen potential cases. These hotlines have struggled with the volume of callers, leading to wait times of hours or, even, an inability to contact health authorities. Symptoma is a symptom-to-disease digital health assistant that can differentiate more than 20,000 diseases with an accuracy of more than 90%. We tested the accuracy of Symptoma to identify COVID-19 using a set of diverse clinical cases combined with case reports of COVID-19. We showed that Symptoma can accurately distinguish COVID-19 in 96.32% of clinical cases. When considering only COVID-19 symptoms and risk factors, Symptoma identified 100% of those infected when presented with only three signs. Lastly, we showed that Symptoma’s accuracy far exceeds that of simple “yes-no” questionnaires widely available online. In summary, Symptoma provides unparalleled accuracy in systematically identifying cases of COVID-19 while also considering over 20,000 other diseases. Furthermore, Symptoma allows free text input, furthered with disease-specific follow up questions, in 36 languages. Combined, these results and accessibility give Symptoma the potential to be a key tool in the global fight against COVID-19. The Symptoma predictor is freely available online at https://www.symptoma.com.

## Introduction

Currently, the world is facing an unprecedented health crisis caused by the novel coronavirus disease (COVID-19). To track and hopefully curb this pandemic, large-scale laboratory testing for COVID-19 is becoming widespread. However, capacities are far from being able to test whole populations. Therefore many countries have established phone hotlines to pre-screen persons who are unsure about their COVID-19 infection status. Only after talking to an operator and being identified as a potential case, often due to being symptomatic, will laboratory testing occur. However, these hotlines are severely overrun worldwide, leading to hour-long waiting periods and, even, disconnected lines. This ultimately leads to many COVID-19 cases going undiagnosed.

One solution to the overwhelming number of calls inundating hotlines is to pre-screen them using computer-based approaches. Digital services are already utilised for assessment and triage in various countries to reduce stress on the emergency responders, e.g., the “digtal 111” service provided by the NHS in England^1, 2^. These methods can be grouped into two categories. Firstly, a large number of simple yes/no online questionnaires are available. These questionnaires lead straight to the point but are limited in their informative value. They do not provide a deep understanding of a patient’s health situation, they do not allow for the consideration of additional symptoms, they do not allow the generation of additional data for analysis, and, critically, they are often language- and/or country-specific.

The alternative to these questionnaires is general-purpose symptom checkers which allow a user to list or select their symptoms before being provided with potential diagnoses. Several have already been developed over recent years (benchmarked in Nateqi *et al.*)^3^. However, most of these symptom checkers are highly restricted in the number of diseases taken into account as building up the underlying databases is fundamentally cost-intensive and slow. Furthermore, language ambiguities, such as synonyms, are hard to overcome, e.g., dyspnea is the medical term for shortness of breath, though is rarely used outside of medical profession. This inevitably leads to small disease databases where users can only choose from a limited list of pre-defined symptoms, lowering their viability when a new disease emerges.

Recently, we showed that Symptoma, a symptom-to-disease digital health assistant, significantly outperforms other symptom checkers in general diagnosis^3^. This was also confirmed by an independent study^4^. In the following, we present the accuracy of Symptoma with regards to systematically identifying cases of COVID-19 while also considering over 20,000 other diseases.

## Results

### Sensitivity and specificity

Symptoma classifies nearly all 30 COVID-19 case descriptions correctly as COVID-19 cases (96.6% sensitivity), failing only when presented with a case containing no defining symptoms of COVID-19 (Case 3006: Fever, Fatigue, Dizziness, Constipation, Rhonchi, Tachypnea, and Bilateral pneumonia). Achieving 100% sensitivity is however easy e.g. by constructing a test that simply classifies every case as COVID-19. To address this issue we also tested how well Symptoma performs on cases of non-COVID-19 patients. For this purpose, we use a set of 1,112 British Medical Journal (BMJ) cases that stretch over 84 fields of medicine (see Methods). Of these 1,112 cases, only 41 are classified as potential COVID-19 cases by Symptoma, with only seven of these ranking COVID-19 higher than the correct diagnoses. These seven cases relate to diseases that present similarly to COVID-19, however, have far lower incidence rates and, therefore, are deemed less likely, e.g. Severe Acute Respiratory Syndrome (SARS-CoV) or the Avian influenza A (H5N1) virus infection (bird flu). The results are summarized in Table 1.

**Table 1.** Sensitivity and specificity of Symptoma in COVID-19 cases and BMJ negative controls.

|  | Flagged as<br>COVID-19 Risk | Not flagged as<br>COVID-19 Risk |
| --- | --- | --- |
| COVID-19 cases<br>(n=30) | 29 (TP) | 1 (FN) |
| BMJ cases<br>(n=1112) | 41 (FP) | 1071 (TN) |
| Sensitivity | 96.66% (29 of 30 detected) |  |
| Specificity | 96.31% (41 of 1112 incorrectly detected) |  |
| Accuracy | 96.32% (1100 of 1142 predictions correct) |  |

### Discovery speed and sensitivity

Identifying patients presenting with COVID-19 both quickly and efficiently is of utmost importance to digital diagnoses. However, achieving both speed and accuracy simultaneously is remarkably difficult. Short, and therefore quick, questionnaires will typically have low specificity, while conversely, long questionnaires lack efficiency and speed, often containing many questions not pertinent to any given patient. Symptoma’s free text search allows quick, efficient, and complex queries of symptom’s without constraint to a predefined list of symptoms.

To highlight this with regards to COVID-19, we show in Figure 1, the search rank of queries containing various numbers of symptoms known to be present in those infected with COVID-19 (see Methods). Key symptoms, such as suffering a fever or dry cough, leads to COVID-19 suggested within the top 30 search results immediately. Living in an area with a high incidence of COVID-19 or contact with a known case of COVID-19 results in a high-risk assessment immediately. The top 30 threshold is passed by 75%, 98.5%, and 100% of one, two, and three symptom queries respectively. At three symptoms, 99.1% of the possible combinations are returned within the top 10 results, and with four symptoms, all queries return COVID-19 within the top 10. These results highlight the speed with which a correct diagnosis can be obtained, even when minimal symptoms are entered into the query.

**Figure 1.**
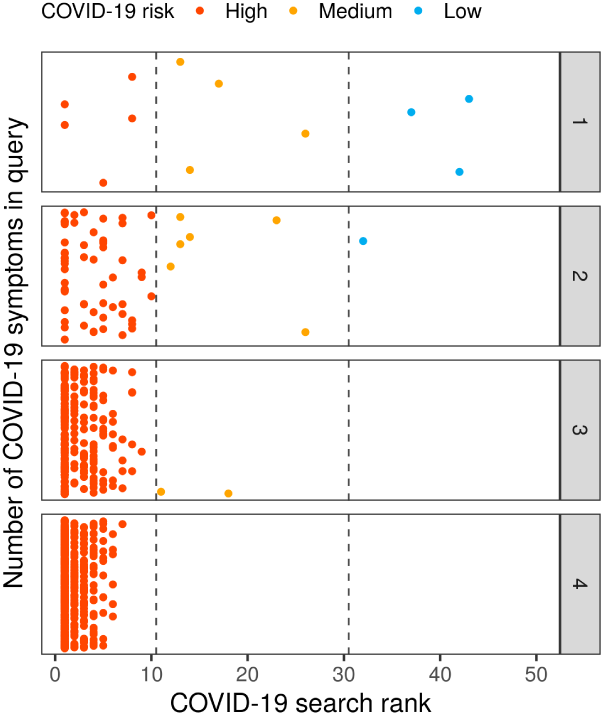
Identification of COVID-19 cases with regards to the number of query terms entered. On the x-axis, the search rank of the query in Symptoma is given against the y-axis where each panel considers a different number of symptoms in the query. All combinations of the reported COVID-19 symptoms are considered with each dot representing one unique combination. Points are jittered vertically for clarity only.

### Symptoma performs better than simple approaches

Below we compare Symptoma to alternative approaches used to predict the likelihood of bearing COVID-19. Using a restricted set of symptoms as input, the probability that a user is suffering from COVID-19 in comparison to either influenza, the common cold or hay fever is calculated based on their respective literature-derived symptom frequencies (see Table S1). We implemented four different methods: the distance in symptom-space between case presentation and symptom frequency (SF-DIST), the distance normalised by the standard deviation (SF-SD), the distance normalised by the first principal component (SF-PCA) and the cosine similarity (SF-COS). These methods are described in detail within the Methods.

To evaluate the performance of these approaches in comparison to Symptoma, we classified the combined COVID-19 and BMJ cases from Table 1, subsetting to only those cases that have at least one COVID-19 symptom (n=406). A case is classified as COVID-19 positive if the probability of COVID-19 higher than the probability for influenza, common cold or hay fever. As Symptoma weights COVID-19 against more than 20,000 other diseases, to provide a fair comparison, we note only the returned rank of influenza, the common cold and hay fever. If COVID-19 is returned first, we classify that case as COVID-19 positive.

The results, summarised in Figure 2a, show that Symptoma performs considerably better than any other approach. A margin of 0.09 sensitivity and 0.20 specificity was found between Symptoma and the other methods, clearly showing the ability of Symptoma to differentiate between COVID-19 and other common causes with similar symptoms. Additionally, we show Symptoma’s performance using our default measurements of accuracy outlined above (see Table 1). On this subset of difficult cases, Symptoma outperforms all other methods, which, is remarkable, as the other approaches consider only four causes as a potential diagnosis.

**Figure 2.**
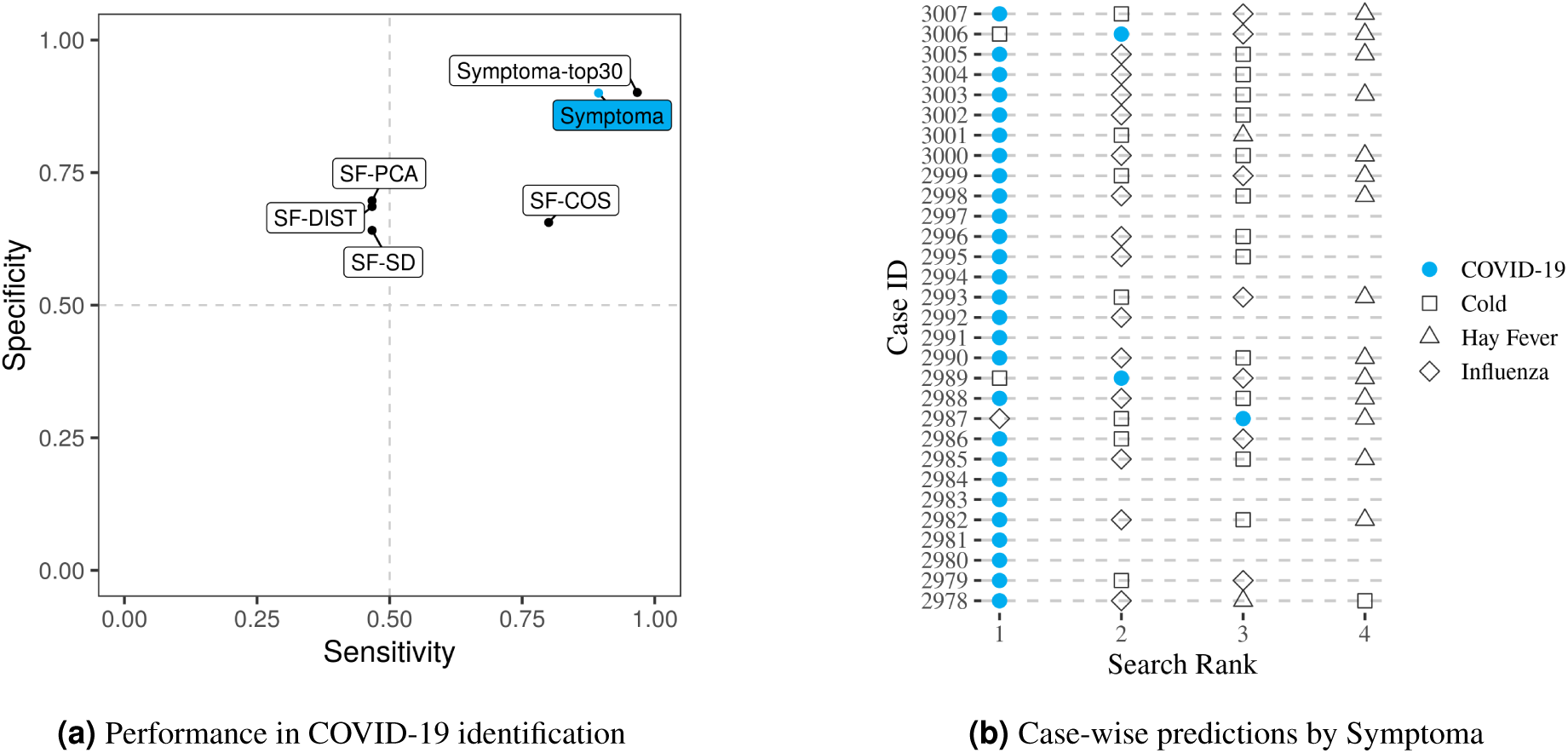
Performance by Symptoma and alternative approaches concerning the identification of COVID-19 cases. On the left, we show the performance of Symptoma, highlighted in blue, against alternatives, all of which are constrained to consider only three alternative diagnoses (the common cold, influenza, and hay fever). We also give Symptoma’s accuracy on this set of case reports when unconstrained (labelled as top30). On the right, we breakdown the predictions by Symptoma on the COVID-19 cases. Missing points indicate that the corresponding disease was considered so unlikely that it was not returned by the search.

In Figure 2b, we breakdown Symptoma’s performance in the above test across our COVID-19 cases. We find that for the three cases that did not report COVID-19 as diagnosis, one is case 3006 outlined above as having no defining features of COVID-19. Case 2889, diagnosed as a common cold, symptoms include Conjunctivitis and Chills, two features more commonly associated with a cold. Case 2987, symptoms include Nasal Congestion and Rhinorrhea, both indications of an alternative diagnosis to COVID-19.

## Discussion

Above, we compared the symptom-to-disease search engine Symptoma to alternative methods on both COVID-19 and BMJ-derived decoy cases. We showed that Symptoma is superior to these alternative approaches in multiple aspects. First, Symptoma can correctly identify cases of COVID-19, not only against a few diseases but against more than 20,000 other potential causes. This far exceeds the 6,000 differential diagnoses provided by the second-largest symptom checker “Isabel Healthcare”^5^. Secondly, we showed that Symptoma needs only a few symptoms to highlight cases of COVID-19. Crucially, these can be inputted as free text that is semantically understood. For example, if one enters “Tiramisu”, “Food Poisoning” is returned as one of the top results. In contrast, other COVID-19 questionnaires and symptom checkers require that patients only select from a predefined list of symptoms or are constrained to fixed questions. Furthermore, Symptoma is localised into 36 languages, allowing a standardised approach on disease predictions globally. Lastly, we found that Symptoma performs better than simple questionnaire-based approached in identifying COVID-19 cases, even though these alternative approaches only considered three alternative diagnoses (COVID-19, influenza, common cold, and hay fever). Moreover, as Symptoma is constantly mining the newest literature, it keeps up-to-date with the latest knowledge and alters its predictions accordingly, e.g. the recent reports of anosmia in COVID-19 patients^6, 7^. Similarly, by tracking cases of COVID-19 entered into Symptoma, due to the free-text function, new symptoms related to COVID-19 may be highlighted. On these grounds, we believe that Symptoma is a highly valuable tool in the global COVID-19 crisis.

## Methods

### Test cases

To show the performance of Symptoma for COVID-19 we analysed a total of 1142 medical test cases. The different sets and sources of these cases are described below.

#### BMJ Cases

A total of 1112 cases were sourced from the British Medical Journal (BMJ) and transcribed by a medical clinician into sets of symptoms, both negative and positive, alongside other risk factors, the patient’s age, and the patient’s sex when available^8, 9^. The cases cover a diverse range of causes, including patients suffering rib fractures, rabies, or metastatic cancer. The number of symptoms and keywords per case ranges from one to 33 (median eight) including complex terms such as “right true vocal cord is immobile”.

#### Covid-19 Cases

A set of 30 case reports were derived from the current literature, the sources of which are listed in full within the Supplementary Information. For each case, a list of symptoms and risk factors the patient presented with is given, alongside their age and sex where available.

#### COVID-19 Cases: computer generated

We make use of the World Health Organisation (WHO) COVID-19 symptom list to construct example queries from those infected with COVID-19^10^. The ten most frequent symptoms are combined with both “contact with someone infected with COVID-19” and “visiting/living in a COVID-19 risk area”, to give 12 possible symptoms and/or risk factors. All possible combinations of these are then taken as potential COVID-19 cases yielding a total number of 4096 artificial cases.

### Accuracy Measurements

For any given set of symptoms, many possible causes could give rise to that specific presentation. We count a prediction as a true positive if the true cause is listed within the top 30 results returned by Symptoma. Note that this is the maximum number of causes returned by Symptoma for any given query. Given the possible 20,000 causes contained within Symptoma, this is the top 0.15%. Focussing on COVID-19, we can generate the following classification:

- True positive: COVID-19 case and COVID-19 returned in top 30 results
- False positive: Non-COVID-19 case and COVID-19 returned in top 30 results
- True negative: Non-COVID-19 case and COVID-19 not returned in top 30 results
- False negative: COVID-19 case and COVID-19 not returned in top 30 results

Throughout, we also assess a more strict threshold of COVID-19 being returned in the top 10 results. We refer to this stringent threshold as the “high-risk” boundary.

### Alternative predictors

We developed four alternative methods to give the likelihood that a given patient has either COVID-19, influenza, common cold or hay fever. Underlying each were the frequencies with which various symptoms presented for these diseases (Table S1). To determine the probability of each disease, we represented each patient case in a 10-dimensional space, where each axis represents a different symptom. A value of one corresponds to exhibiting the symptom, zero means the patient does not have the symptom, and 0.5 means that they are unsure. This is fundamentally equivalent to many of the simple questionnaire-based approached to COVID-19 identification.

In the most simplistic approach (SF-DIST), we calculated the distance in space between the patient and each of the four diseases, each of which can also be seen as a point in the 10-dimensional symptom space. Normalisation yields the respective probabilities. In the second approach, the same procedure is used, but the distance in each dimension is scaled by the respective standard deviation of each symptom across all diseases (SF-SD). In the third approach, the distance in each dimension is scaled by the first principal component of a matrix consisting of all symptoms across all diseases (SF-PCA). Lastly, we interpreted the points as vectors and calculated the cosine similarity between the cases and diseases (SF-COS).

For assessing the accuracy of these approaches, we used the following criteria

- True positive: COVID-19 case and COVID-19 returned as most probable cause
- False positive: Non-COVID-19 case and COVID-19 returned as most probable cause
- True negative: Non-COVID-19 case and COVID-19 not returned as most probable cause
- False negative: COVID-19 case and COVID-19 not returned as most probable cause

## Contributors

Study design: BK, AM, JN.

Data compilation: SG, NM, IA.

Data analysis: AM, BK, NM, IA.

Writing the manuscript: BK, AM.

Revising the manuscript critically: AM, BK, JN, SG.

## Data sharing

Results can be freely reproduced using the web interface https://www.symptoma.com/

## Acknowledgments

This study has received funding from the European Union’s Horizon 2020 research and innovation programme under grant agreement No 830017.

