## Supplemental Information for "An artificial intelligence-based first-line defence against COVID-19: digitally screening citizens for risks via a chatbot"

### Supplementary Information

#### Symptom Frequencies

| | COVID-19 | Common cold | Influenza | Hay fever |
|---|---|---|---|---|
| Fever | 87.9 [1] | 15 [3] | 68 [6] | – |
| Fatigue | 38.1 [1] | 42 [4] | 94 [6] | – |
| Dry cough | 67.7 [1] | 80 [3] | 93 [6] | 22 [10] |
| Sneezing | – | 74 [4] | 58 [7] | 96 [11] |
| Malaise | 14.8 [1] | 30 [4] | 94 [6] | – |
| Rhinorrhea | 4 [2] | 95 [3] | 91 [6] | 62.1 [12] |
| Sore throat | 13.9 [1] | 70 [3] | 84 [6] | 30 [10] |
| Diarrhea | 3.7 [1] | 11 [4] | 14.4 [8] | – |
| Headache | 13.6 [1] | 80 [5] | 91 [6] | 50 [13] |
| Dyspnea | 18.6 [1] | 21 [4] | 63 [9] | – |

**Table 1.** Symptom frequencies as extracted from the literature.

[1] https://www.who.int/docs/default-source/coronaviruse/who-china-joint-mission-on-covid-19-final-report.pdf
[2] https://www.thelancet.com/action/showPdf?pii=S0140-6736%2820%2930211-7
[3] https://www.sciencedirect.com/science/article/pii/S0095454305703559?via%3Dihub
[4] https://www.ncbi.nlm.nih.gov/pubmed/3057962
[5] https://www.ncbi.nlm.nih.gov/pmc/articles/PMC4347877/pdf/nihms658637.pdf
[6] https://jamanetwork.com/journals/jamainternalmedicine/fullarticle/485554
[7] https://www.ncbi.nlm.nih.gov/pmc/articles/PMC4915903/
[8] https://www.ncbi.nlm.nih.gov/pmc/articles/PMC4676820/
[9] https://www.ncbi.nlm.nih.gov/pmc/articles/PMC3650195/
[10] https://www.ncbi.nlm.nih.gov/pubmed/10971479
[11] https://www.ncbi.nlm.nih.gov/pmc/articles/PMC5806744/
[12] https://www.researchgate.net/publication/307953143_Inverse_correlation_of_soluble_programmed_cell_death-1_ligand-1_sPD-L1_with_eosinophil_count_and_clinical_severity_in_allergic_rhinitis_patients
[13] https://www.ncbi.nlm.nih.gov/pubmed/17300360

### COVID-19 Case Reports

| Case ID | Source |
| --- | --- |
| 3007 | Zhang X, Song W, Liu X, Lyu L. CT image of novel coronavirus pneumonia: a case report. Jpn J Radiol. 2020 Mar 18. doi: 10.1007/s11604-020-00945-1. |
| 3006, 2987, 2986, 2985 | Wang Z, Chen X, Lu Y, Chen F, Zhang W. Clinical characteristics and therapeutic procedure for four cases with 2019 novel coronavirus pneumonia receiving combined Chinese and Western medicine treatment. Biosci Trends. 2020 Mar 16;14(1):64-68. doi: 10.5582/bst.2020.01030. |
| 3005 | Thevarajan I, Nguyen THO, Koutsakos M, et al. Breadth of concomitant immune responses prior to patient recovery: a case report of non-severe COVID-19. Nat Med. 2020. doi: 10.1038/s41591-020-0819-2 |
| 3004 | Ng K, Poon BH, Kiat Puar TH, et al. COVID-19 and the Risk to Health Care Workers: A Case Report. Ann Intern Med. 2020. doi: 10.7326/L20-0175 |
| 3003, 3002 | Chen L, Liu W, Zhang Q, et al. RNA based mNGS approach identifies a novel human coronavirus from two individual pneumonia cases in 2019 Wuhan outbreak. Emerg Microbes Infect. 2020 Feb 5;9(1):313-319. doi: 10.1080/22221751.2020.1725399 |
| 3001, 2997 | Lin X, Gong Z, Xiao Z, Xiong J, Fan B, Liu J. Novel Coronavirus Pneumonia Outbreak in 2019: Computed Tomographic Findings in Two Cases. Korean J Radiol. 2020 Mar;21(3):365-368. doi: 10.3348/kjr.2020.0078 |
| 3000 | Lim J, Jeon S, Shin HY, et al. Case of the Index Patient Who Caused Tertiary Transmission of COVID-19 Infection in Korea: the Application of Lopinavir/Ritonavir for the Treatment of COVID-19 Infected Pneumonia Monitored by Quantitative RT-PCR. J Korean Med Sci. 2020 Feb 17;35(6):e79. doi: 10.3346/jkms.2020.35.e79. |
| 2999 | Park WB, Kwon NJ, Choi SJ et al. Virus Isolation from the First Patient with SARS-CoV-2 in Korea. J Korean Med Sci. 2020 Feb 24;35(7):e84. doi: 10.3346/jkms.2020.35.e84. |
| 2998 | An P, Song P, Lian K, Wang Y. CT Manifestations of Novel Coronavirus Pneumonia: A Case Report. Balkan Med J. 2020 Mar 6. doi: 10.4274/balkanmedj.galenos.2020.2020.2.15. |
| 2996 | Wei J, Xu H, Xiong J, et al. 2019 Novel Coronavirus (COVID-19) Pneumonia: Serial Computed Tomography Findings. Korean J Radiol. 2020 Apr;21(4):501-504. doi: 10.3348/kjr.2020.0112. |
| 2995 | Zhu L, Xu X, Ma K, et al. Successful recovery of COVID-19 pneumonia in a renal transplant recipient with long-term immunosuppression. Am J Transplant. 2020 Mar 17. doi: 10.1111/ajt.15869. |
| 2994 | Wang S, Guo L, Chen L, et al. A case report of neonatal COVID-19 infection in China. Clin Infect Dis. 2020 Mar 12. pii: ciaa225. doi: 10.1093/cid/ciaa225. |
| 2993 | Chen D, Xu W, Lei Z, et al. Recurrence of positive SARS-CoV-2 RNA in COVID-19: A case report. Int J Infect Dis. 2020 Mar 5. pii: S1201-9712(20)30122-3. doi: 10.1016/j.ijid.2020.03.003. |
| 2992 | Xu Z, Shi L, Wang Y, et al. Pathological findings of COVID-19 associated with acute respiratory distress syndrome. Lancet Respir Med. 2020 Feb 18. pii: S2213-2600(20)30076-X. doi: 10.1016/S2213-2600(20)30076-X. |
| 2991 | Kim JY, Ko JH, Kim Y, et al. Viral Load Kinetics of SARS-CoV-2 Infection in First Two Patients in Korea. J Korean Med Sci. 2020 Feb 24;35(7):e86. doi: 10.3346/jkms.2020.35.e86. |
| 2990, 2989, 2988 | Bernard Stoecklin S, Rolland P, Silue Y, et al. First cases of coronavirus disease 2019 (COVID-19) in France: surveillance, investigations and control measures, January 2020. Euro Surveill. 2020 Feb;25(6). doi: 10.2807/1560-7917.ES.2020.25.6.2000094. |
| 2984 | Shi H, Han X, Zheng C. Evolution of CT Manifestations in a Patient Recovered from 2019 Novel Coronavirus (2019-nCoV) Pneumonia in Wuhan, China. Radiology. 2020 Apr;295(1):20. doi: 10.1148/radiol.2020200269. |
| 2983, 2982 | Fang Y, Zhang H, Xu Y, Xie J, Pang P, Ji W. CT Manifestations of Two Cases of 2019 Novel Coronavirus (2019-nCoV) Pneumonia. Radiology. 2020 Apr;295(1):208-209. doi: 10.1148/radiol.2020200280. |
| 2981 | Liu P, Tan XZ. 2019 Novel Coronavirus (2019-nCoV) Pneumonia. Radiology. 2020 Apr;295(1):19. doi: 10.1148/radiol.2020200257. |
| 2980 | Lei J, Li J, Li X, Qi X. CT Imaging of the 2019 Novel Coronavirus (2019-nCoV) Pneumonia. Radiology. 2020 Apr;295(1):18. doi: 10.1148/radiol.2020200236. |
| 2979 | Park JY, Han MS, Park KU, Kim JY, Choi EH. First Pediatric Case of Coronavirus Disease 2019 in Korea. J Korean Med Sci. 2020 Mar 23;35(11):e124. doi: 10.3346/jkms.2020.35.e124. |
| 2978 | Hosoda T, Sakamoto M, Shimizu H, Okabe N. SARS-CoV-2 enterocolitis with persisting to excrete the virus for about two weeks after recovering from diarrhea: A case report. Infect Control Hosp Epidemiol. 2020 Mar 19:1-4. doi: 10.1017/ice.2020.87. |
